# Antiviral immune responses, cellular metabolism and adhesion are differentially modulated by SARS-CoV-2 ORF7a or ORF7b

**DOI:** 10.1101/2022.06.01.494101

**Authors:** Tránsito García-García, Raúl Fernández-Rodríguez, Natalia Redondo, Ana de Lucas-Rius, Sara Zaldívar-López, Blanca Dies López-Ayllón, José M. Suárez-Cárdenas, Ángeles Jiménez-Marín, María Montoya, Juan J. Garrido

**Affiliations:** Immunogenomics and Molecular Pathogenesis BIO365 Group, Department of Genetics, University of Córdoba, Córdoba, Spain; Maimónides Biomedical Research Institute of Córdoba (IMIBIC), GA-14 Research Group, Córdoba, Spain; Molecular Biomedicine Department, Centro de Investigaciones Biológicas Margarita Salas (CIB), CSIC, Madrid 28040, Spain

**Keywords:** SARS-COV-2, ORF7a, ORF7b, COVID-19

## Abstract

SARS-CoV-2, the causative agent of the present COVID-19 pandemic, possesses eleven accessory proteins encoded in its genome, and some have been implicated in facilitating infection and pathogenesis through their interaction with cellular components. Among these proteins, accessory protein ORF7a and ORF7b functions are poorly understood. In this study, A549 cells were transduced to express ORF7a and ORF7b, respectively, to explore more in depth the role of each accessory protein in the pathological manifestation leading to COVID-19. Bioinformatic analysis and integration of transcriptome results identified defined canonical pathways and functional groupings revealing that after expression of ORF7a or ORF7b, the lung cells are potentially altered to create conditions more favorable for SARS-CoV-2, by inhibiting the IFN-I response, increasing proinflammatory cytokines release, and altering cell metabolic activity and adhesion. Based on these results, it is reasonable to suggest that ORF7a and ORF7b could be targeted by new therapies or used as future biomarkers during this pandemic.

## Introduction

There is an urgent need to better understand the molecular mechanisms governing severe acute respiratory syndrome coronavirus 2 (SARS-CoV-2), the causative agent of the ongoing coronavirus disease 2019 (COVID-19) pandemic. SARS-CoV-2 belongs to the family *Coronaviridae*, subfamily *Orthocoronavirinae*, genus *Betacoronavirus*, subgenus *Sarbecovirus*(Gorbalenya et al., 2020). Since the 2019/2020 outbreak, SARS-CoV-2 has infected more than 500 million people, causing more than 6 million deaths worldwide (https://covid19.who.int/), and unleashing a serious global health problem. COVID-19 exhibits a broad spectrum of severity and progression patterns from mild upper respiratory disease or even asymptomatic sub-clinical infection to severe and fatal pneumonia (Rajarshi et al., 2021).

SARS-CoV-2 genome consists of a single-stranded positive-sense RNA of 29,903 bp containing 14 open reading frames (ORFs) encoding 31 viral proteins (Ellis et al., 2021). Although much of the research on this virus is focused on the Spike protein (Walls et al., 2020; Yang et al., 2020; Zost et al., 2020), recent reports demonstrate that SARS-CoV-2 accessory proteins are involved in COVID-19 pathogenesis by modulating antiviral host responses (Jiang et al., 2020; Konno et al., 2020; Miorin et al., 2020; Wang et al., 2021; Wu et al., 2021; Xia et al., 2020; Zhang et al., 2021). Eleven accessory proteins are encoded in the SARS-CoV-2 genome (Redondo et al., 2021) and some of them have been involved in facilitating the infection process by interacting with cell components (Gordon et al., 2020; Stukalov et al., 2021). Among these proteins, accessory protein ORF7a and ORF7b are less studied and their functions have not been fully resolved. ORF7a is a type-I transmembrane protein of 121 amino acid residues with an N-terminal signal peptide (residues 1–15), an Immunoglobulin (Ig)-like ectodomain (residues 16–96), a transmembrane domain (residues 97–116), and an endoplasmic reticulum (ER) retention motif (residues 117–121) (Zhou et al., 2021) (**Figure 1a**). It exhibits 95.9% sequence similarity with ORF7a protein from SARS-CoV (Yoshimoto, 2020). SARS-CoV-2 ORF7a Ig-like ectodomain has recently been identified as an immunomodulating factor able to interact with CD14^+^ monocytes, leading to a decrease in their antigen-presenting ability and triggering a dramatic inflammatory response (Zhou et al., 2021). In addition, ORF7a is one of SARS-CoV-2 proteins able to antagonize IFN-I responses (Xia et al., 2020) by promoting inhibition of IFN-I signaling *via* STAT2 phosphorylation (Cao et al., 2021). ORF7b is a 43 amino acid transmembrane protein, one less than in SARS-CoV (**Figure 1a**). Although less well studied than ORF7a, some authors identified that ORF7b is able to multimerize through a leucine zipper and hypothesized that it could interfere with some cellular processes involving leucine zipper formation and epithelial cell-cell adhesion that might underlie some common COVID-19 symptoms such as heart rate dysregulation and odor loss (Fogeron et al., 2021). Recently, ORF7b has been involved in host immune responses by promoting IFN-β, TNF-α and IL-6 expression, activating IFN-I signaling pathways and eventually accelerating TNF-induced apoptosis (Yang et al., 2021). As there is still a lack of knowledge regarding the role of ORF7a and ORF7b in the pathogenesis of SARS-CoV-2, the response of A549 human epithelial cells expressing ORF7a or ORF7b separately was analyzed using transcriptomic approaches combined with bioinformatic analysis and functional assays. Overexpression of ORF7a or ORF7b induced specific and differential alteration on metabolic cascades via UGT1A9, PTGS2 and CYP1A1; interferon responses via OASL, IFIT1 and IFIT2; inflammation via IL-8, IL-11 and CXCL1 and cell adhesion via ICAM-1, ZO-1 and *γ*-catenin. Overall, we found that the expression of either ORF7a or ORF7b was sufficient to alter cellular networks in a manner similar to full SARS-CoV-2 virus infection.

**Figure 1.**
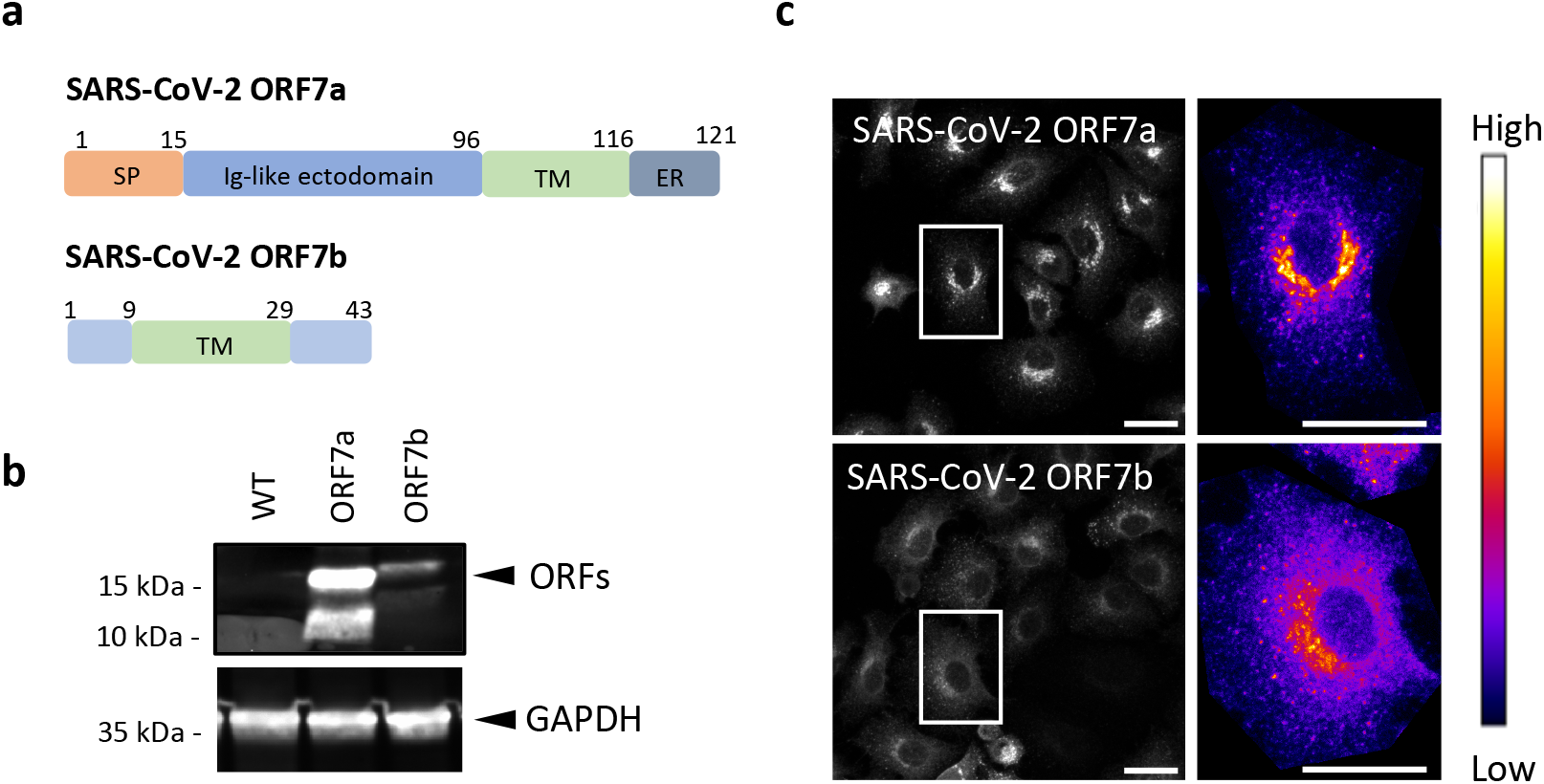
Expression of SARS-CoV-2 ORF7a and ORF7b in A459 epithelial cells. **a**, Schematic representation of SARS-CoV-2 ORF7a and ORF7b proteins. Domains are highlighted in couleur (SP, signal peptide; TM, transmembrane domain; ER, endoplasmic reticulum retention signal) and numbers denote the residues sites. **b**, Expression of SARS-CoV-2 proteins ORF7a and ORF7b. C-terminally Strep-tagged viral proteins were transduced in A459 cells and analyzed by western blotting using anti-Strep-tag and anti-GAPDH antibodies. **c**, Cellular localization of ORF7a and ORF7b. A459 transduced cells with Strep-tagged SARS-CoV-2 proteins were imaged by confocal microscopy. Right panels are heatmaps of signal intensity detected in a representative cell. Scale bar is 25 µm.

## Results

### ORF7a and ORF7b overexpression in A549 cells

SARS-CoV-2 uses several strategies to interact with and interfere with the host cellular machinery. To investigate the function of ORF7a and ORF7b in such interactions, A549 human lung carcinoma cell line was used since infection of lung epithelial cells is a hallmark of SARS-CoV-2 infection in the humans. Lentiviruses expressing individual viral proteins ORF7a or ORF7b were transduced in A549 cells with a 2xStrep-tag at C-terminus to allow their detection. In western blot analysis of A549-ORF7a and A549-ORF7b cells, proteins bands of 15 kDa were detected using an ant-Strep-tag antibody in agreement with previous results (Xia et al., 2020) (**Figure 1b**). For ORF7a, an additional band of 10 kDa was also detected, which may due to protein cleavage. Protein overexpression was confirmed by immunofluorescence and highlighted different patterns of localization in A549 cells. According to a heatmaps of signal intensities in a single cell, ORF7a is highly concentrated in the perinuclear region while ORF7b was diffused through the cytoplasm with an enrichment adjacent to the nucleus (**Figure 1c**). Because protein localization can provide important information on their function, immunofluorescence confocal microscopy was assessed to examine the subcellular co-localization of proteins with different organelles. The A549 cells were stained with anti-Strep-tag and the organelle markers GM130 to visualize Golgi, Tom20 to visualize Mitochondria and Rab4 and Rab7 to visualize early and late Endosomes, respectively. Cytosolic ORF7a C-terminal contains a di-lysine ER retrieval signal (KRKTE) that mediates protein trafficking to the ER-Golgi intermediate compartment (**Figure 1a**). Thus, ORF7a localized predominantly at the Golgi apparatus (**Figure 2a**) in agreement with previous studies (Gordon et al., 2020; Lee et al., 2021; Zhang et al., 2020) and with early Endosomes. Likewise, ORF7b has been associated with ER in SARS-CoV-2 (Lee et al., 2021; Zhang et al., 2020) and with Golgi in SARS-CoV (Schaecher et al., 2007, 2008). However, our results showed that SARS-CoV-2 ORF7b partially localizes with Golgi (Pearson’s coefficient of 0,58) and predominantly colocalized with Mitochondria (Pearson’s coefficient of 0,74) **(Figure 2b**).

**Figure 2.**
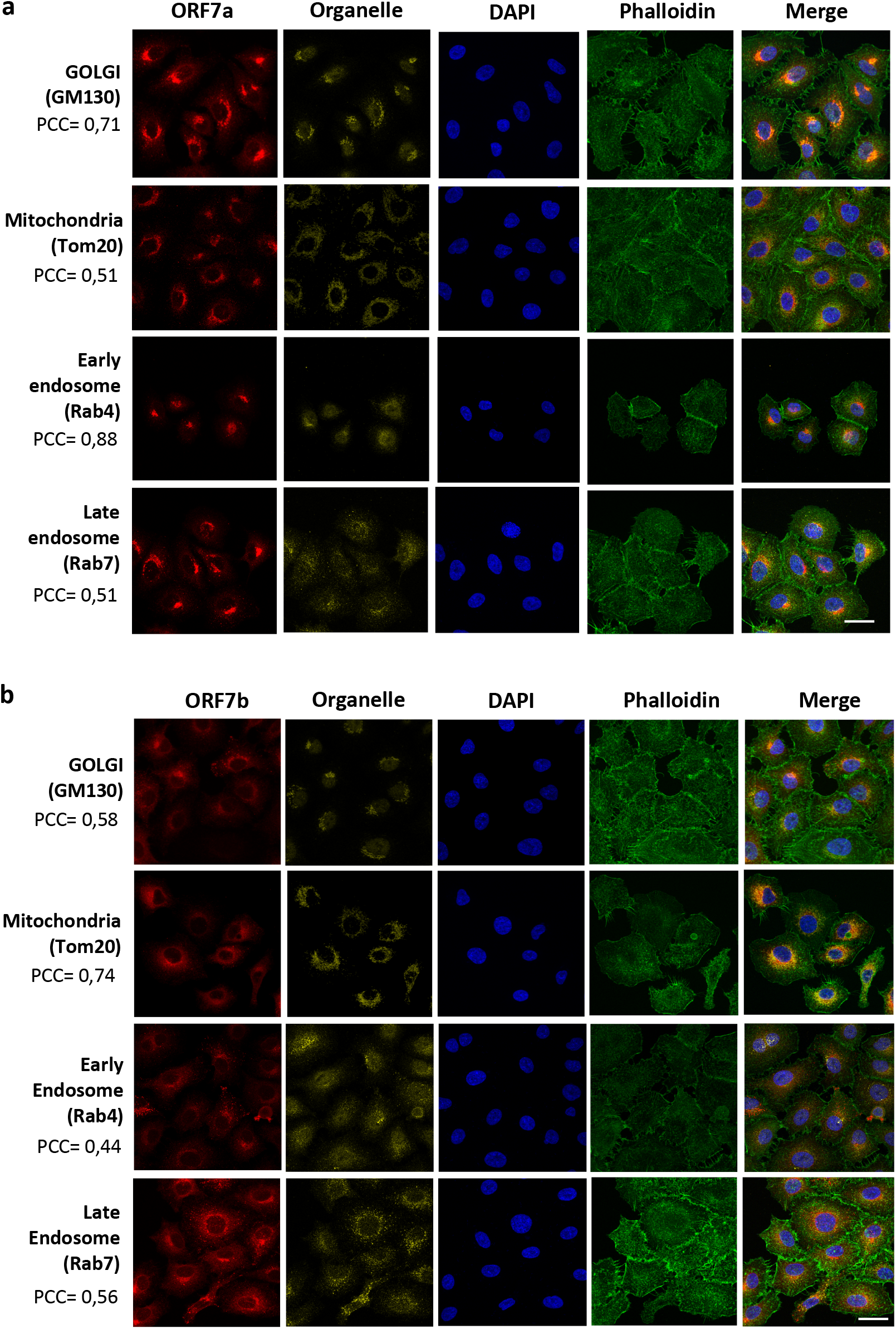
Cellular localization of SARS-CoV-2 ORF7a and ORF7b. Confocal analysis of SARS-CoV-2 protein ORF7a and ORF7b localization in A549 cells transduced with Strep-tagged-ORF7a (**a**) or ORF7b (**b**) and organelle markers: GM130 (Golgi), Tom20 (Mitochondria) Rab4 (Early endosome) and Rab7 (Late endosome). Red: Strep-tag antibody signal; yellow: organelle markers; Blue: DAPI (nuclei staining); Green: Phalloidin. Scale bar, 25 μm. All experiments were done at least twice, and one representative is shown. PCC indicates the Pearson’s coefficient for co-localization of each organelle with the Strep-tag ORF7a or ORF7b.

### RNA sequencing identified genes altered on A549-ORF7a and A549-ORF7b cells

Differential gene expression analysis was performed for A549 cells and A549 cells expressing either ORF7a or ORF7a (**Figure S1a**). A principal component analysis (PCA) based on normalized counts from DESeq2 was used to explore the similarity of our samples. High quality was achieved since samples were well clustered (**Figure S1b**). We identified the overall up- and down-regulated differentially expressed genes (DEGs) in A549-ORF7a and A549-ORF7b cells compared to A549 non-transduced cells (adjusted *p*-value <0,05 and log2 fold chance >1). In total, 882 genes were up-regulated, and 457 genes were down-regulated by ORF7a (**Figure 3a**, Table S1). Likewise, 652 genes were up-regulated and 500 genes were down-regulated in A549-ORF7b cells (**Figure 3a**, Table S2). To delineate the potential functions of SARS-CoV-2 ORF7a and ORF7b proteins, gene ontology (GO) and pathways (KEGG) analysis was conducted based on their respective DEGs. The most significantly enriched biological processes in cells expressing ORF7a were *cell-cell adhesion, extracellular structure* and *extracellular matrix organization* (**Figure S2a**, Table S1) and pathways such as *steroid hormone biosynthesis, ascorbate and aldarate metabolism, bile secretion* and *ECM-receptor interaction* (**Figure S2c**, Table S1). In A549-ORF7b expressing cells, processes like *extracellular structure, extracellular matrix organization* and *metabolism pathways* were also the most significant ones (**Figure S2b and S2d**, Table S2). These data indicated that identified DEGs are mainly enriched in ECM-related items and metabolism genes, suggesting that both proteins could be interacting in similar pathways. The overlap of DEGs in ORF7a- and ORF7b-regulated genes was analyzed using the Venn diagram visualization, showing 751 genes in common of which 311 genes were up-regulated and 440 genes were down-regulated (**Figure 3b**). Next, the MCODE enrichment analysis based on this PPI network was applied to the common genes. Out of the eight PPI modules presented, three of them were discarded since they contained only 3 genes (**Figure 3c**, Table S3). Among the top list of enriched terms of MCODE 1 (22 genes), three major Reactome pathways were *extracellular matrix, integrin cell surface interactions* and *collagen chain trimerization*, most of which were down-regulated (red and green colors). The top list of MCODE 2 enriched categories (15 genes) included GO-term *positive regulation of vasculature development, positive regulation of angiogenesis* and the reactome pathway *adherens junctions interactions (JUP, CDH7, CDH5, PDZD3* down-regulated and *CDH6* up-regulated). MCODE 3 (11 genes) was associated with *post-translational protein phosphorylation, regulation of Insulin-like Growth Factor (IGF) transport and uptake by Insulin-like Growth Factor Binding Proteins (IGFBPs)* and *NABA CORE MATRISOME* pathways, most of them up-regulated (blue and violet colors). MCODE 4 (10 genes) included the KEGG pathway *steroid hormone biosynthesis* and the *metapathway biotransformation Phase I and II*. Finally, MCODE 5 (5 genes: *WNT5A, IL7R, VAMP8, STON1, SYT1*) was associated with the Reactome pathway *Cargo recognition for clathrin-mediated endocytosis, Clathrin-mediated endocytosis* and *Membrane Trafficking* (**Figure 3c**).

**Figure 3.**
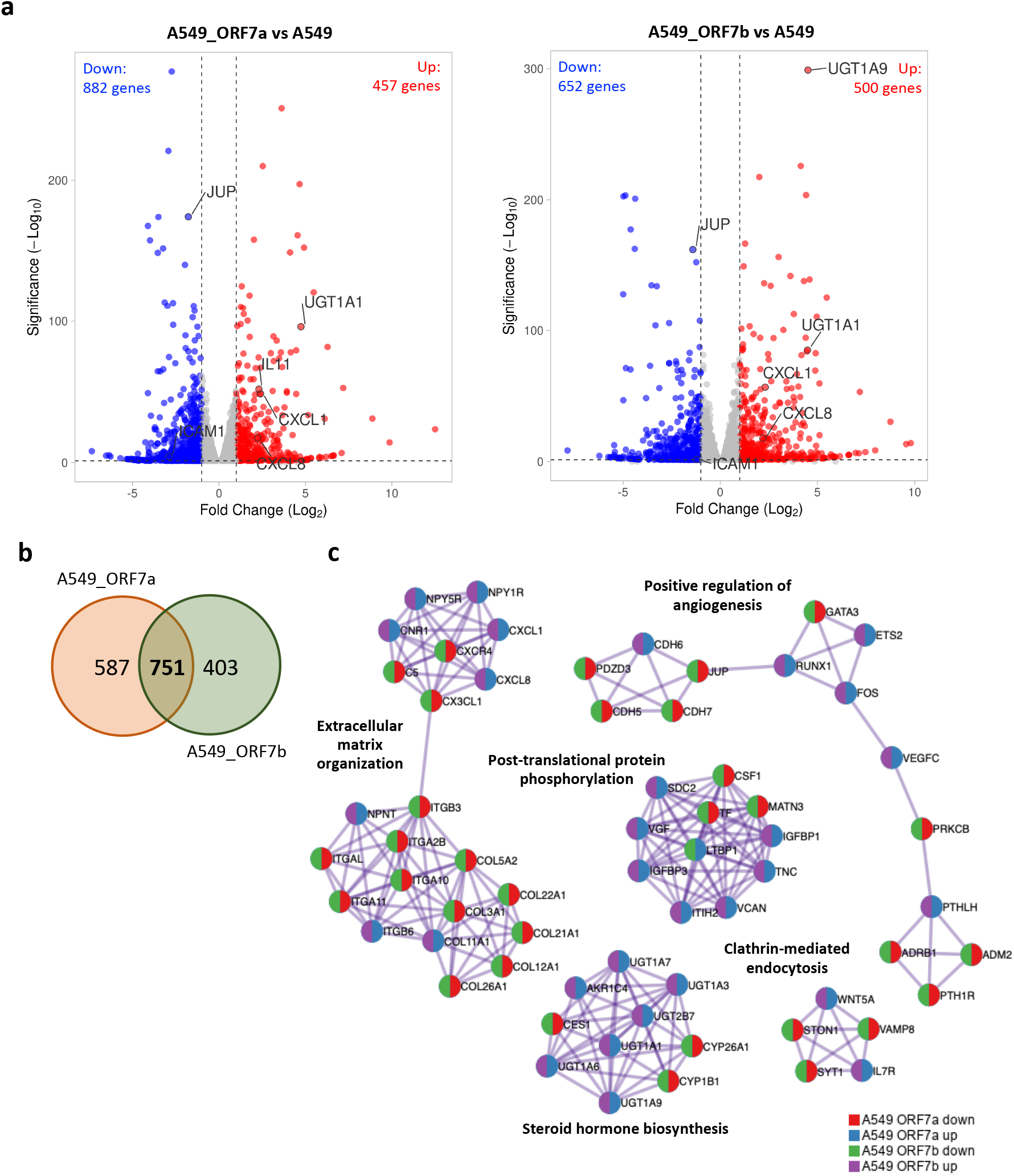
a, Identification of DEGs in A549 cells expressing ORF7a or ORF7b. **a**, Volcano plot of DEGs in A549 cells transduced with SARS-CoV-2 ORF7a (*left*) or ORF7b (*right*) compared with A549 WT. **b**, Venn diagram of DEGs in A549 cells expressing ORF7a and ORF7b. **c**, MCODE enrichment analysis by Metascape. MCODE algorithm was applied to clustered enrichment ontology terms to identify neighborhoods where proteins are densely connected and GO enrichment analysis was applied to each MCODE network to assign biological meanings. The color code for pie sector represents a gene list.

### Expression of ORF7a and ORF7b induced metabolic disfunctions in A549 cells

The uridin diphosphate (UDP)-glucuronosyltransferase (UGT) family of enzymes catalyzes the attachment of a glucuronic acid (glucuronidation) to certain drugs and xenobiotics, as well as to endogenous compounds such as bilirubin to facilitate their elimination from the body (Mano et al., 2018). We found genes coding for several UDP-glucuronosyltransferases (UGT1A1, UGT1A3, UGT1A6, UGT1A7, UGT1A9, UGT2B7) highly overexpressed in transduced cells with ORF7a and ORF7b (**Figure 3c and 4a**). Using qRT-PCR, the expression of accessory proteins ORF7a and ORF7b induced an 18-fold increase in *UGT1A9* mRNA levels as compared with control cells (**Figure 4b**).

**Figure 4.**
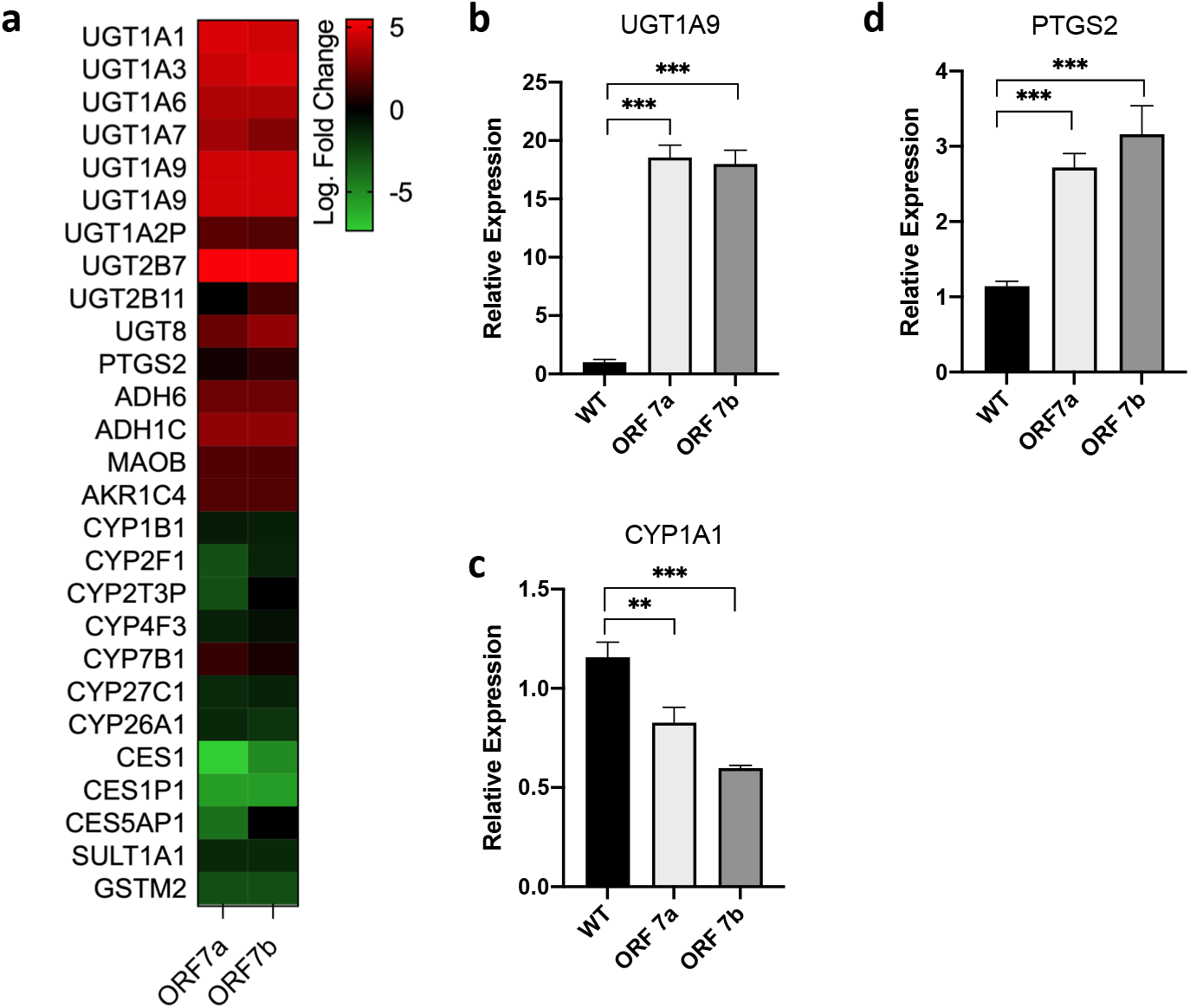
SARS-CoV-2 ORF7a and ORF7b alter the metabolic process. **a**, Log2 Foldchange Heatmaps of DEGs involved in metabolism pathways. **b-c**, Expression of UGT1A9 (b) and CYP1A1 (c) genes were calculated with 2^-ΔΔCT^ method by normalizing to that of GADPH. The fold changes were calculated with respect to the level of A549 WT. Error bars represent mean ± SD (n=3). Statistical significance is as follows: \*\**p*□<□0.01, \*\*\**p*□<□0.001.

On the other hand, cytochromes P450 (P450s or CYPs) comprise a superfamily of monooxygenase enzymes that catalyze oxygen insertion into a large array of different substrates such as lipids and steroids, as well as drug compounds and other xenobiotics. A prominent member of the P450 superfamily is CYP1A1, which metabolizes a variety of substrates including fatty acids such as arachidonic acid, the fluoroquinolone antibiotic difloxacin, and the drug theophylline. CYP1A1 also acts on several procarcinogenic molecules (Munro, 2018). Thus, CYP’s system is the most important drug-metabolizing enzyme family existing amongst species(Stipp and Acco, 2021). Generally, CYPs’ expression or activity decrease during viral infections, chronic inflammation and in presence of proinflammatory cytokines (e.g. IL-6, TNF-α, IFN-γ TGF-β and IL-1β), which can result in alterations of the pharmacological effects of substances in inflammatory diseases (Christmas, 2015; Wang et al., 2022). Surprisingly, ORF7a and ORF7b overexpression led to significant down-regulation of genes coding for CYP enzymes such as CYP1B1, CYP2F1, CYP2T3P, CYP4F3, CYP27C1 and CYP26A1 (**Figure 4a**). We also observed a reduction in *CYP1A1* expression by qRT-PCR in A549-ORF7a and A549-ORF7b cells (**Figure 4c**). These results indicate that both ORF7a and ORF7b perturb the UGT and CYP expression suggesting a metabolic dysfunction that might facilitate the metabolic inactivation of drug therapies in COVID-19 patients.

Interestingly, we found that the gene coding for the prostaglandin-endoperoxidase synthase 2 (PTGS2), also known as cyclooxygenase 2 (COX-2) was significantly up-regulated in A549-ORF7a and ORF7b cells (**Figure 4d**). COX-2 enzyme is critical for the generation of prostaglandins, lipid molecules with diverse roles in maintaining homeostasis as well as in mediating pathogenic mechanisms, including the inflammatory response (Ricciotti and FitzGerald, 2011). Stimulation of PTSG2 has been previously detected in SARS-CoV via its spike and nucleocapsid proteins (Liu et al., 2007; Yan et al., 2006). More recently it was found that SARS-CoV-2 induced COX-2 upregulation in human lung epithelial cells (Blanco-Melo et al., 2020) as observed in the present study.

### ORF7a and ORF7b disrupt inflammatory responses balance in A549 cells

SARS-CoV-2 infection induces unbalanced inflammatory responses, characterized by weak production of type I interferon (IFN-I) and overexpression of proinflammatory cytokines, both of which are linked to severe clinical outcomes (Hojyo et al., 2020; Sa Ribero et al., 2020). Data from our RNAseq study showed a down-regulation of several Interferon-Stimulated Genes (ISGs) as well as an up-regulation of cytokines and chemokines (**Figure 5a**). As previously reported, SARS-CoV-2 ORF7a antagonizes the production of IFN-I by blocking the phosphorylation of STAT2 thereby suppressing the transcriptional activation of antiviral ISGs (Martin-Sancho et al., 2021; Xia et al., 2020). As expected, we found reduced expression of *OASL, IFIT1, IFIT2* and *TRIM22* genes in A549-ORF7a cells (**Figure 5a**). However, in ORF7b-expressing cells we observed a decrease of *IFIT1* and *TRIM22* expression bur overexpression of *IFITM1* and *RSAD2* genes, suggesting that both proteins may interfere with IFN-I responses but employing different strategies. To confirm this, we examined by qRT-PCR whether ORF7a or ORF7b expression affected ISG-associated gene expression in A549 cells. The results showed that while ORF7a inhibited the expression of *OASL, IFIT1* and *IFIT2*, ORF7b did not (**Figure 5b**), indicating that ORF7a is an IFN-I antagonist whereas ORF7b is not.

**Figure 5.**
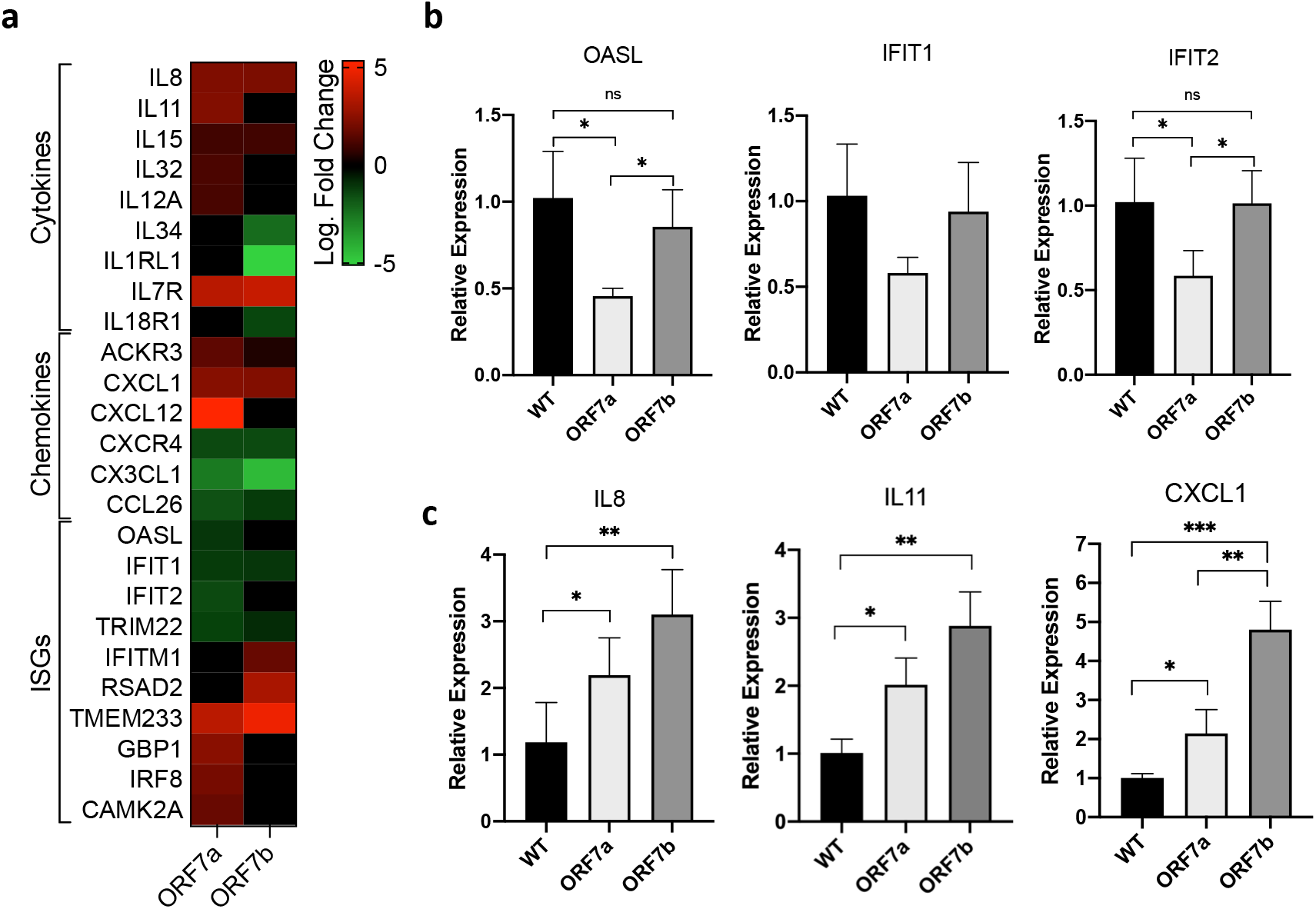
Interferon and inflammatory responses to SARS-CoV-2 ORF7a and ORF7b. **a**, Heatmaps showing expression of genes related to antiviral and inflammatory response compared to A549 cells. Genes shown in red are significantly increased, genes in green significantly decreased and genes in black indicate no change in expression. **b-c**, Expression of ISGs, OASL, IFIT1 and IFIT2 (**b**) and cytokines IL8, IL11 and CXCL1 (**c**) were calculated with 2^-ΔΔCT^ method by normalizing to that of GADPH. The fold changes were calculated with respect to the level of A549 WT. Error bars represent mean ± SD (n=3). Statistical significance is as follows: \**p*□<□0.05, \*\**p*□<□0.01, \*\*\**p*□<□0.001.

ORF7a is also thought to activate the NFkB pathway and promote the production of inflammatory cytokines, which play a significant role in the clinical severity of COVID-19 (Su et al., 2021). Conversely, little is known about ORF7b’s role in the inflammatory response. It has been described that ORF7b may promote the IFN-I signaling pathways and eventually accelerate TNF-induced apoptosis (Yang et al., 2021). An increase in several genes encoding cytokines was observed here, including *IL-8, IL-11, IL-15, IL-32* and *IL-12A* in A549-ORF7a and *IL-8* and *IL-15* in A549-ORF7b (**Figure 5a**). In addition, genes encoding chemokines such as ACKR3, CXCL1 and CXCL12 were strongly overexpressed in ORF7a cells whereas only *CXCL1* was observed overexpressed in ORF7b cells. To further corroborate this data, expression of *IL-8, IL-11* and *CXCL1* was measured in A549-ORF7a and A549-ORF7b by qRT-PCR. Increased levels of these genes were detected in both transduced cell lines **(Figure 5c**). All in all, our results showed that accessory proteins ORF7a and ORF7b play key roles in regulating the host immune responses to SARS-CoV-2 infection, inhibiting the IFN-I production by ORF7a and increasing proinflammatory cytokines release by both ORF7a and ORF7b.

### ORF7a and ORF7b expression alter cell-ECM and cell-cell interactions

Most of the dysregulated genes associated with the enriched process *ECM organization* (**Figure 3c**) were down-regulated. Among them, we found genes coding for integrins (*ITGA2B, ITGB3, ITGB6, ITGA10, ITGA11*), collagens (*COL4A4, COL9A3*), tenascins (*TNC*), vitronectin (*VTN*), laminins (*LAMB2, LAMA3, LAMB3, LAMC3*), nephronectin (*NPNT*) and thrombospondin-3 (*THBS3-AS1*) and cell adhesion molecules (CAMs) such as immunoglobulin superfamily members (ICAM-1, IGSF11), integrins (ITGAL), cadherins (CDH5) and claudins (CLDN2), most of them down-regulated (**Figure 6a**). Integrins are integral cell-surface proteins composed of an alpha chain and beta chain that participate in cell adhesion as well as in cell-surface mediated signaling. Interestingly, *ITGA2B* and *ITGB3* genes encoding alpha and beta chains of the alpha-IIb/beta-3 integrin were observed down-regulated. This integrin is highly expressed in platelet and plays a crucial role in the blood coagulation system by mediating platelet aggregation (Ma et al., 2007). Integrin alpha-IIb/beta-3 binds specific adhesive proteins as fibrinogen/fibrin, plasminogen, prothrombin, thrombospondin and vitronectin (Huang et al., 2019; Ma et al., 2007). A down-regulation of *ITGA10* and *ITGA1*, the genes encoding the beta chains of the collagen-binding integrins alpha-10/beta-1 and the alpha-11/beta-1, respectively (White et al., 2004), were also observed. Conversely, we observed an increase in the expression of the gene encoding beta-6 integrin subunit (ITGB6), which forms exclusively a dimer with the alpha v chain (alpha-v/beta-6) (Bandyopadhyay and Raghavan, 2009). This integrin plays a role in modulating the innate immune response in lungs, being able to bind ligands such as fibronectin and transforming growth factor beta-1 (TGF-1). Integrin alpha-v/beta-6 is predominantly expressed in epithelial cells and is highly expressed in inflamed and injured lung tissue (Horan et al., 2008). In summary, our results suggest that ORF7a and ORF7b expression is responsible for critical changes in the expression of important components of the cell adhesion machinery, which can significantly alter cell–ECM and cell-cell interactions. To test this hypothesis, transduced and control cells were evaluated for their adhesion ability to different ECM components (**Figure 6b**). According to transcriptomic data, A549-ORF7a cells showed weaker binding capacity to fibrinogen, collagen IV, and fibronectin whereas A549-ORF7b exhibited lower adhesion to fibrinogen and fibronectin. This could be due to the inhibition of *ITGA2B, ITGB3, ITGA10* and *ITGA11* expression observed in both transduced cells.

**Figure 6.**
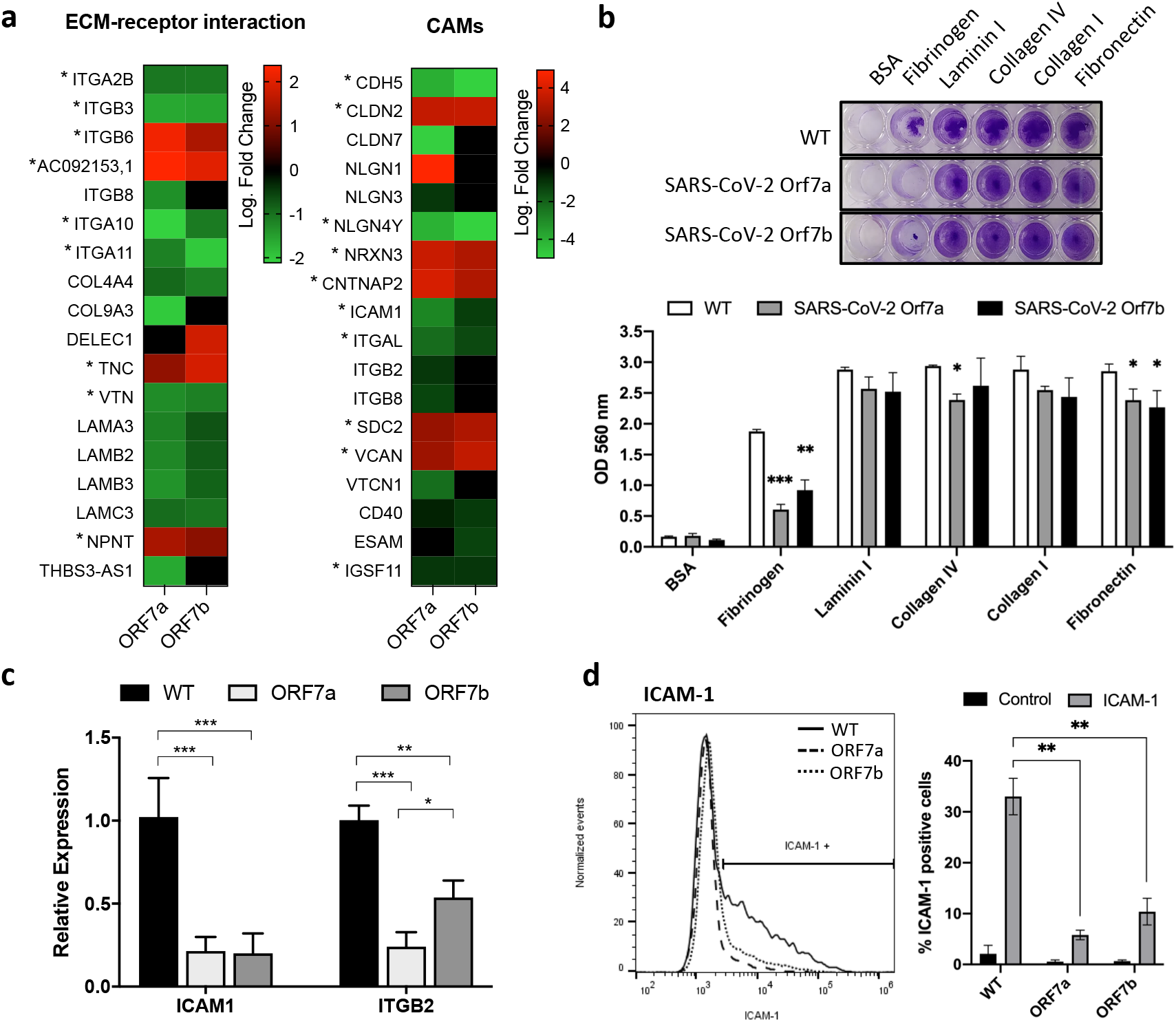
Effect of SARS-CoV-2 ORF7a and SARS-CoV-2 ORF7b in cell adhesion. **a**, Log2 Foldchange Heatmaps of DEGs involved in ECM-receptor interaction (hsa04512) and cell adhesion molecules (CAMS) (hsa04514). Red for up-regulated, green for down-regulated and asterisk (*) for significant in both cell lines. **b**, Quantification of A549 cells adhering to the extracellular matrix (ECM) components fibronectin, collagen I, collagen IV, laminin I, and fibrinogen, and BSA (control). **c**, RT-qPCR for ICAM1 and ITGB2 genes. **d**, Analysis by flow cytometry of ICAM-1 expression in A549 cells expressing ORF7a or ORF7b (*left panel*) and quantification of the percentage of ICAM-1 positive cells (*right panel*). Data are represented as mean□±□SD (n=3). Statistical significance is given as follows: \**p*□<□0.05, \*\**p*□<□0.01, \*\*\**p*□<□0.001 to the control group A549 WT.

Integrins typically bind to the ECM while immunoglobulin members and cadherins are associated with cell adhesion and cell-cell signaling. In fact, previous studies have suggested that SARS-CoV-2 ORF7a may play a vital role in recognizing macromolecules on the cellular surface since its structural homology with the human intercellular adhesion family of molecules (ICAM) (Nizamudeen et al., 2021; Zhou et al., 2021). Interaction between ICAM and integrins regulates immune cell migration, activation, and target cell recognition, which are important host defense mechanisms against infectious agents. Interestingly, our transcriptomic results revealed a down-regulation of *ICAM1* as well as *ITGAL* and *ITGB2*, the genes encoding the integrin alpha-L and beta-2 chains for the leukocyte function-associated antigen-1 (LFA-1) (**Figure 6a**). Significant reductions in *ICAM1* and *ITGB2* expression were further confirmed by qRT-PCR in A549-ORF7a and A549-ORF7b cells (**Figure 6c**). Next, flow cytometry was used to analyze ICAM-1 expression on cell surfaces. As shown in **Figure 6d**, ICAM-1 expression was significantly reduced in direct relation to ORF7a and ORF7b expression in A549 cells.

Cadherins are the major cell adhesion molecules (CAMs) responsible for Ca2+-dependent cell-cell adhesion and they are therefore crucial for promoting diverse morphogenetic processes (Hirano and Takeichi, 2012). The transcriptomic analyses of A549-ORF7a and ORF7b revealed a reduction in transcripts encoding adherens junctions proteins including cadherins (*CDH5, CDH7, CDH19*), protocadherins (*PCDHGA5, PCDHGB5, PCDH10, PCDH17*), and catenins such as γ-catenin (*JUP*, also known as plakoglobin). Additionally, dysregulation of genes encoding tight junction and gap junction proteins such as claudin 2 (*CLDN2*), zona occludens protein 2 (*TJP2*) and the gap junction protein beta 3 (*GJB3*) were also observed (**Figure 7a**). To assess the impact of these transcriptional changes on cell-cell junctions in the bronchoalveolar epithelium, plakoglobin expression in transduced A549 cells was analyzed by Western blotting. Plakoglobin is present in desmosomes and adherens junctions being critical for desmosome components recruitment which are required for junction assembly (Sumigray and Lechler, 2015). As expected, plakoglobin expression in A549-ORF7a and ORF7b cells was firstly analyzed by western blotting. As expected, A549-ORF7a and A549-ORF7b cells had significantly reduced compared to A549 cells, with ORF7b having a more pronounced effect (**Figure 7b**). Next, a cell-cell adhesion assay was performed in order to detect potential defects in cell-cell adhesive bonds. Interestingly, we detected a reduced adherence associated to ORF7b but not to ORF7a (**Figure 7c**), probably linked to γ-catenin reduction. To better characterize the barrier function of the A549 transduced cells, immunofluorescence analysis in the tight junction protein zonula occludens-1 (ZO-1) was performed. ZO-1 is a scaffold protein that promotes protein-protein complex assembly and influences the structure and function of lung epithelial barrier (Bazzoni et al., 2000). After 8 days of culture on transwell inserts to polarized the cells, ZO-1 was localized in membranes of neighboring epithelial cells showing continuous distribution in A549 cells and A549-ORF7a while fluorescence intensity decreased and distribution was discontinuous in membranes from A549-ORF7b cells (**Figure 7d**). In agreement with this result, we noticed a greater facility to trypsinize A549-ORF7b when we cultured them (data not shown). Together, these data show that ORF7b could disrupt cell to cell interaction leading to a breakdown of the lung epithelial barrier.

**Figure 7.**
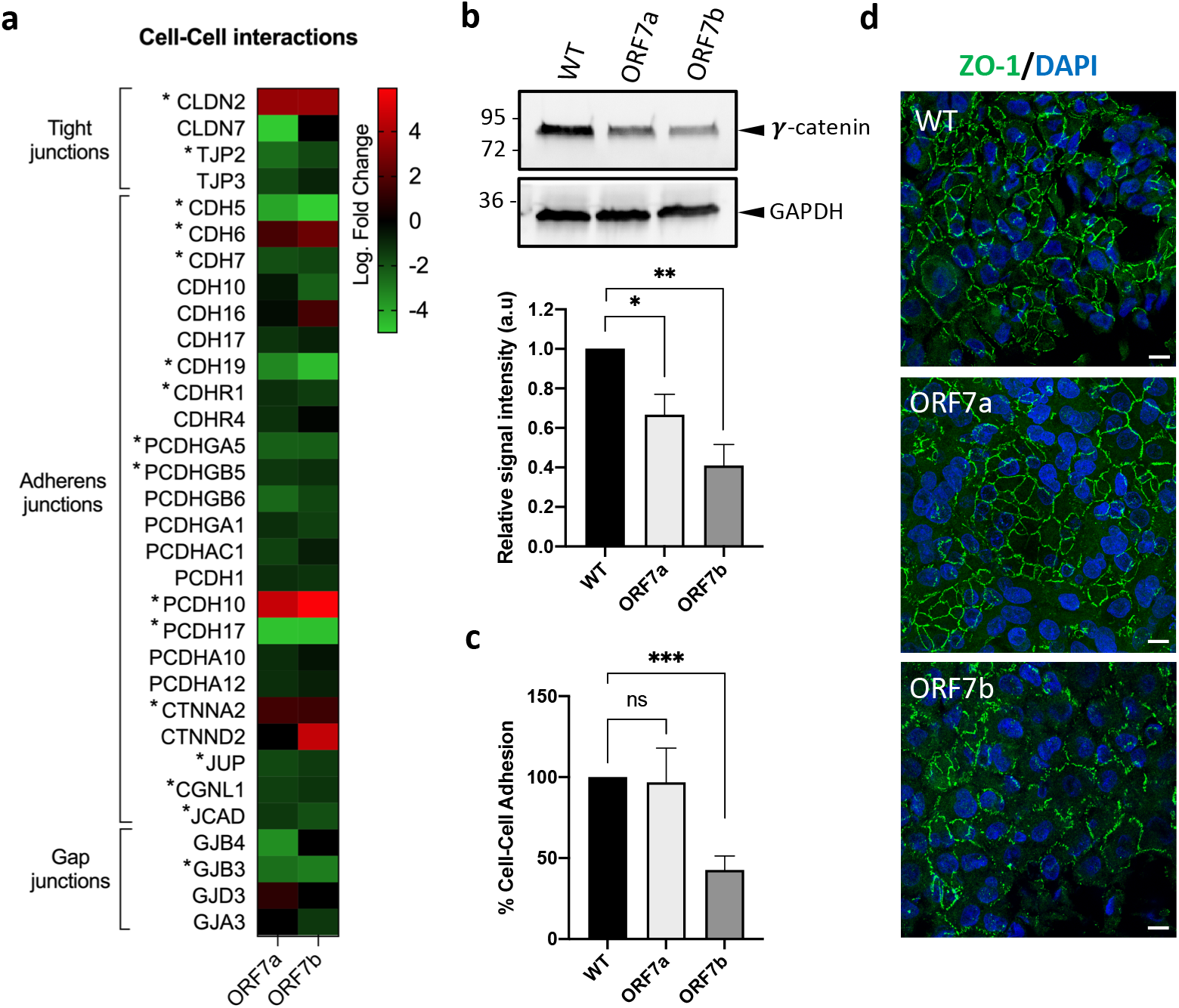
Effect of SARS-CoV-2 ORF7a and SARS-CoV-2 ORF7b in cell junctions. **a**, Log2 Foldchange Heatmaps of DEGs related to cell-cell adhesion interaction. **b**, Western blotting showing the expression of Y-catenin, the major protein in cell-cell adhesion at the desmosomes (*top panel*) and quantification for Y-catenin (*bottom panel*). GAPDH was used as a control. **c**, Cell-cell adhesion assay showing the percentage of adherent cells. Data are represented as mean□±□SD (n=3). Statistical significance is given as follows: \**p*□<□0.05, \*\**p*□<□0.01, \*\*\**p*□<□0.001 to the control group A549 WT. **d**, Immunostaining for the tight junction protein ZO-1 after 8 days of culture on transwell inserts. Scale bar, 10 μm.

### Host transcriptome responses by ORF7a and ORF7b resembled responses to SARS-CoV-2 infection

A comparative study was performed by integrating transcriptomic results from SARS-CoV-2 infections deposited in public repositories and those generated in the present work. The objective was to analyze the presence of common alteration in common genes, in order to estimate which aspects of the host response to SARS-CoV-2 infection could be attributed to the expression of the accessory proteins ORF7a and ORF7b. To this end, we compared our differential expression data with those obtained from infecting ACE2-transfected A549 cells with SARS-CoV-2 and others resulting from transcriptomic analysis of post-mortem lung biopsies from COVID-19 patients (GSE147507) (Blanco-Melo et al., 2020) (**Figure S3a-b**). Several ORF7a and ORF7b-associated genes were found to be differentially expressed upon SARS-CoV-2 infection in A549-ACE2 cells and in clinical COVID-19 lung samples (**Figure 8a-b**). These data suggested a certain degree of similarity between transcriptional changes associated with ORF7a and ORF7b expression and SARS-CoV-2 infection.

**Figure 8.**
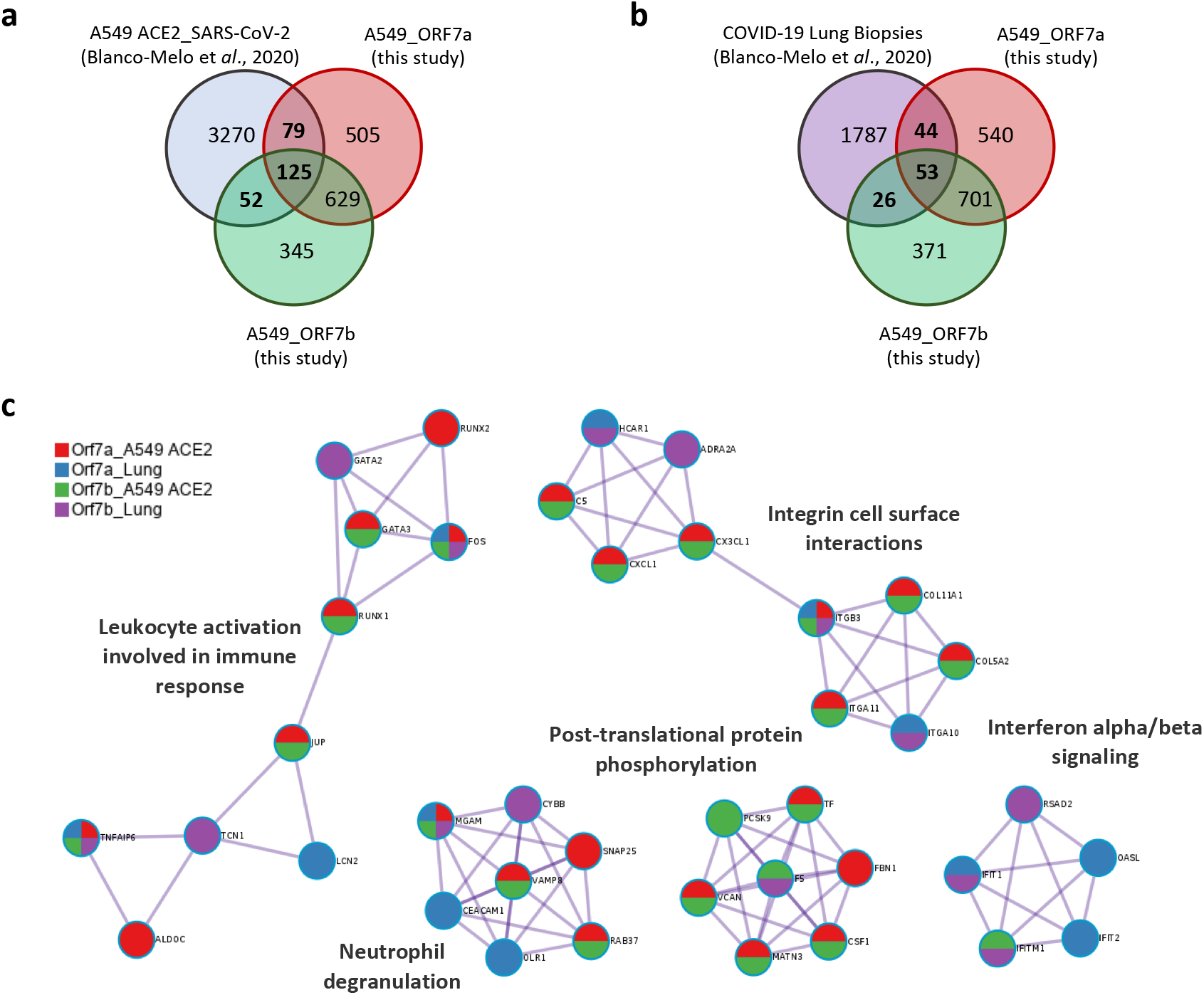
Comparison of host transcriptome responses of ORF7a and ORF7b to SARS-CoV-2 infection. **a-b**, Venn diagrams of the intersection between differentially expressed genes in A549 cells expressing ORF7a or ORF7b and SARS-CoV-2-regulated genes in A549-ACE2 cells (a) and COVID-19 lung biopsies (b). Data are from Blanco-Melo et *al*., 2020. **c**, MCODE components identified from the PPI network. Network nodes are displayed as pies. The color code for the pie sector represents a gene list (metascape.org).

Totally, 204 DEG in A549-ORF7a and 177 in A549-ORF7b were identified also differentially expressed in SARS-CoV-2 infected A549-ACE2 cells. Of these, 125 genes were regulated by both accessory proteins (**Figure 8a**). When compared with data sets from post-mortem lung biopsies, 97 genes were found associated with ORF7a, 79 genes with ORF7b and 53 genes were in common (**Figure 8b**). To further investigate differential perturbation of pathways regulated by these accessory proteins in SARS-CoV-2 infection, GO-enrichment analysis and PPI network were contained with to obtain MCODE network components (**Figure 8c**, Table S4). Overall, substantially enrichment biological process of common genes belonging to the inflammatory response (GO006954) (**Figure S3c**) and enrichment pathways of the MCODE network, such as, *leukocyte activation involved in immune response, neutrophil degranulation, integrin cell surface interactions, post-translational protein phosphorylation* and *interferon alpha/beta signaling* were detected. Among *leukocyte activation involved in immune response* included RUNX1, RUNX2, GATA2, GATA3, FOS, JUP, LCN2, TCN1, ALDOC and TNFAIP6, TNF alpha-induced protein 6 (TNFAIP6) and the AP-1 transcription factor subunit (FOS) belonging to the TNFA signaling via NFKB, that were also detected in our set of dysregulated genes in A549-ORF7a and A549-ORF7b cells. Similarly, they were dysregulated in A549-ACE2 cells infected with SARS-CoV-2 (red and green colors, respectively) and in clinical COVID-19 lung samples (blue and violet colors) **(Figure 8c)**. A dysregulation of 7 neutrophil-enriched genes (MGAM, CYPBB, SNAP25, VAMP8, RAB37, CEACAM1 and OLR1) and the maltase-glucoamylase (MGAM), which is directly involved in neutrophil recruitment and activity (Ericson et al., 2014), was found in all data lists **(Figure 8c)**. Another dysregulated gen in infected A549-ACE2 cells with SARS-CoV-2, in COVID-lung samples and in A549-ORF7a and A549-ORF7b was integrin beta 3 subunit (ITGB3). This gene belongs to the MCODE network integrin cell surface interaction that also includes integrins alpha subunits ITGA10 and ITGA11, collagens COL11A1 and COL5A2, and chemokine CX3CL1, which can act as a ligand for the alpha v/beta 3 integrin **(Figure 8c**). Most of these genes were down-regulated **(Figure 3c)**.

## Discussion

SARS-CoV-2 encodes eleven accessory proteins that modulate host immune responses and may contribute to the pathogenicity of the viral infection (Redondo et al., 2021). Among those accessory proteins, SARS-CoV-2 ORF7a and ORF7b functions are not fully understood. Therefore, human bronchoalveolar A549 epithelial cells individually expressing accessory proteins SARS-CoV-2 ORF7a and ORF7b were generated by lentivirus transduction in order to gain deeper knowledge of their function (**Figure 1**). As previously reported, we observed that SARS-CoV-2 ORF7a localizes at the Golgi apparatus (Gordon et al., 2020; Lee et al., 2021; Zhang et al., 2020). In addition, we also detected this protein in the endosome (**Figure 2a**). However, SARS-CoV-2 ORF7b partially localizes with Golgi and predominantly with mitochondria in A549 epithelial cells (**Figure 2b**), in contrast to the previous observation in COS-7 and Caco-2 cells, where SARS-CoV-2 ORF7b colocalizes with Endoplasmic Reticulum (Lee et al., 2021; Zhang et al., 2020).

Using transcriptomic combined with bioinformatic analysis we have found that individually expressed accessory proteins ORF7a and ORF7b were sufficient to alter cellular networks in a manner that resembles full SARS-CoV-2 (**Figure 3** and **Figure 8**). For the first time to our knowledge, our results showed that SARS-CoV-2 ORF7a and ORF7b were individually implicated in multiple deregulations of host pathways, such as immune response, metabolism, extracellular matrix organization and cell adhesion, broadening our understanding of the pathogenicity of SARS-CoV-2 (**Figure 3**). Interestingly, perturbed UGT and CYP gene expression were altered, genes that encode UDP-glucuronosyltransferase and cytochrome P450 enzymes, respectively (**Figure 4**). Both undoubtedly having an important influence on cell responses to endogenous and exogenous factors, that may impact on disease progression as well as on drug responses. Al-Kuraishy et al. suggested that hyperbilirubinemia in patients with Gilbert syndrome attenuates COVID-19-induced metabolic disturbances (Al-Kuraishy et al., 2021). This syndrome is characterized by genetic alteration of the UGT1A1 gene causing reduced gene expression, resulting in a reduction of the UDP-glucuronosyl transferase function, which is responsible for the conjugation of bilirubin. The antioxidant role of hyperbilirubinemia in COVID-19 patients has been also reported by Liu et al. (Liu et al., 2020) and its beneficial role has been additionally highlighted by Khurana et al., who postulated the use of administration of bilirubin to combat systemic dysfunctions associated with COVID-19 (Khurana et al., 2021). In this study, we observed high expression of *UGT1A1* (log2FC>4), together with some other UGT genes such as *UGT1A9* (**Figure 4**) suggesting that ORF7a and ORF7b may induce metabolic disorders which could alter the levels of bilirubin and other metabolites and may facilitate inactivation of drug therapies that could explain the inefficiency of most of the COVID-19 treatments used. Moreover, many drugs used in COVID-19 patients are metabolized by CYPs (Wang et al., 2022); therefore, if CYPs’ expression is greatly altered in COVID-19 patients, it may contribute to altered drug pharmacokinetics and increased drug-related side effects.

COX-2 enzyme has a central role in viral infections and regulates the expression levels of many serum proteins, including proinflammatory cytokines (Capuano et al., 2020; Ricciotti and FitzGerald, 2011). COX-2 hyperproduction has been reported in lung epithelial cells of patients who died of H5N1 infection (Lee et al., 2008) and detected in blood samples of severe COVID-19 patients (Passos et al., 2022), suggesting that this enzyme is associated with poor clinical outcomes in viral infections. In SARS-CoV, transfection of plasmids encoding either the spike or the nucleocapsid genes was sufficient to stimulate COX-2 expression (Liu et al., 2007; Yan et al., 2006). Similarly, ORF7a and ORF7b proteins may mediate COX-2 induction in SARS-CoV-2 (**Figure 4**).

Another factor in virus pathogenicity and transmissibility is to know how SARS-CoV-2 may subvert immune responses by evading innate immunity. IFN is among the first cytokines to be upregulated in virus-infected cells and are important in coordinating the antiviral response and inflammation. Many SARS-CoV-2 accessory proteins are known to antagonize or evade IFN responses, which may contribute to the delayed IFN expression observed in COVID-19 patients. Both SARS-CoV-2 ORF7a and ORF7b have been reported that inhibit IFN-I signaling by blocking phosphorylation of STAT proteins. Specifically, ORF7a only inhibits STAT2 phosphorylation while ORF7b suppresses both STAT1 and STAT2 phosphorylation (Cao et al., 2021; Xia et al., 2020). Consistent with these studies, a dysregulation of the IFN-I signaling was observed here which was characterized by a reduction in several ISGs expression **(Figure 5**); however only ORF7a seemed to be a potent antagonist of IFN-I. What seems clear is that SARS-CoV-2 has evolved ways of using accessory proteins to subvert the IFN response(Martin-Sancho et al., 2021; Xu et al., 2021), although the antagonistic mechanism requires further study.

Finally, diffuse alveolar damage and alveolar-capillary barrier breakdown are prominent pathological features of severe COVID-19 (Batah and Fabro, 2021) and SARS-CoV-2 ORF7a and ORF7b have been suggested to play a determinant role in the disruption on the lung epithelium in COVID-19 patients (Batah and Fabro, 2021). Here, alterations in gene expression were identified in a large number of critical cell–ECM and cell-cell junction proteins, which were strongly repressed (**Figure 6** and **Figure 7)** leading to a weaker association to selective ECM components as fibrinogen, collagen IV, and fibronectin (**Figure 6b**) and lower cell adhesion (**Figure 7c**). Thus, it is tempting to speculate that upon infection, ORF7a and ORF7b participated in COVID-19 pathogenesis by the destruction of the epithelial barrier. These accessory proteins could disrupt adherents and tight junctions which would contribute to the desquamation of the alveolar wall, as observed in lung biopsies from SARS-CoV-infected patients (Nicholls et al., 2003), macaques (Li et al., 2005) and recently in lung autopsy samples from patients with SARS-CoV–2 infection (D’Agnillo et al., 2021). Based on our results, we propose that alteration of cell junctions would create a breach in the epithelial barrier allowing virions to reach the basal matrix, which compromises endothelial function and “blood-air barrier” integrity. In addition to driving severe lung injury, the breach of the blood-air barrier would permit viral entry into the systemic circulation, where the virus can then disseminate to distant organs leading a widespread multi-organ damage (Ackermann et al., 2020; Varga et al., 2020). However, the molecular mechanism that contributes to the destruction of the alveolar walls remains unclear. Several studies suggest that compromised barrier becomes leaky, permitting alveolar fluid accumulation, and the development of pneumonia and inflammatory cell infiltration. The resulting damage triggers a strong production of cytokines and chemokines (“cytokine storm”) that contribute to the massive recruitment of leukocytes at the site of infection and suggests that the inflammatory response further damages pulmonary cells (D’Agnillo et al., 2021; Hanchard et al., 2020). Here, we also identified that SARS-CoV-2 ORF7a and ORF7b strongly induced the expression of pro-inflammatory cytokines such as IL-8, IL-11 and the chemokine CXCL1 (**Figure 5c**), which could enhance epithelial barrier disruption and subsequently contribute to immune cells infiltration. IL-8 and CXCL1 act as chemoattracts for several immune cells, especially neutrophils to the site of infection regulating immune and inflammatory responses (Allen and Kurdowska, 2014; De Filippo et al., 2013). IL-8 has been considered as a biomarker for disease prognosis of COVID-19 since serum IL-8 was easily detectible in COVID-19 patients with mild symptoms, while serum IL-6 became obviously elevated in severe COVID-19 patients (Del Valle et al., 2020; Li et al., 2021, 2020). Recently, a positive feedback loop of systemic and neutrophil autocrine IL-8 production has been identified, which leads to a prothrombotic neutrophil phenotype characterized by degranulation and neutrophil extracellular trap (NET) formation in severe COVID-19 (Kaiser et al., 2021). Consistent with this neutrophil response, differential expression of genes associated with neutrophil degranulation were detected related to the expression of ORF7a and/or ORF7b (**Figure 8c**). In addition, differential expression of genes involved in leukocyte activation was also observed, including genes of the TNFA signaling via NFkB (**Figure 8c**) such as TNFAIP6 and FOS that might contribute to leukocyte infiltration. Additionally, a high expression of IL-11 was observed (**Figure 5c**). IL-11 is a pleiotropic cytokine with effects that overlap with IL-6, particularly relevant in epithelial cells (West, 2019). In human lungs, IL-11 upregulation has been associated with viral infections and a range of fibroinflammatory diseases, including idiopathic pulmonary fibrosis (Ng et al., 2020). Infection with SARS-CoV-2 has also been associated with fibrosis. Notably, the major risk factors for COVID-19 are shared with those of idiopathic pulmonary fibrosis (George et al., 2020).

Taken together, our findings suggest that SARS-CoV-2 accessory proteins ORF7a and ORF7b trigger changes in human pulmonary epithelial cells, which could likely contribute to, at least in part, metabolic disturbances, diffuse alveolar damage and the hyperinflammatory state in severe COVID-19. Given the results shown in this study, it is tempting to speculate that ORF7a and ORF7b could be used as a target for new therapies against COVID-19 to ameliorate their effects on the lungs. Alternatively, they might be useful as biomarkers in this pandemic.

## Materials and methods

### Lentivirus production, cell culture and transduction

ORF7a or ORF7b coding sequences (codon-optimized for mammalian expression) were cloned into pLVX-EF1α-IRES-Puro Cloning and Expression Lentivector (System Biosciences) to generate pseudotyped lentiviral particles encoding the ORF7a or ORF7b accessory proteins of SARS-CoV-2 (Wuhan-Hu-1 isolate) at the CNIC (Centro Nacional de Investigaciones Cardiovasculares) Viral Vector Unit (ViVU). Briefly, VSV-G pseudotyped ORF7a or ORF7b lentivirus was produced by co-transfection of HEK293T cells with the pLVX-ORF7a or pLVX-ORF7b plasmids, pCMV-Gag-Pol and pCMV-VSV-G using Lipofectamine 2000 Reagent (Thermo Fisher Scientific) as per manufacturer instructions. The supernatant was collected 2 days after transfection, centrifuged at 500 x g for 10 min, and concentrated using Amicon Ultra-15 Centrifugal Filter Units, and finally titrated by serial dilution. A549 pulmonary epithelial cells (ATCC CRM-CCL-185) were transduced by incubating with lentivirus at a MOI of 10 for 24 h followed by 2 µg/ml puromycin treatment to start the selection of successfully transduced cells.

### Western blot

Transduced cells were harvested and lysed in ice-cold RIPA lysis buffer containing complete protease inhibitor and phosphatase inhibitors cocktails (Sigma-Aldrich) at 4° C for 30 min. Cell lysates was mixed with SDS-Laemli sample buffer and heated at 95° C for 5 min. Protein samples were resolved by SDS polyacrylamide gel electrophoresis and transferred onto a nitrocellulose membrane using Trans-Blot Turbo Transfer System (Bio-Rad, Hercules, CA), followed by blocking for 1 h with 5% nonfat milk in Tris-buffered saline-Tween 20 buffer and probing with antibodies against Strep Tag (SAB2702215, Sigma-Aldrich), *γ*-catenin (sc-514115, Santa Cruz Biotechnology) and GAPDH (A00084, GenScript) (Supplementary Table 6). The washed membranes were incubated with secondary antibody StarBright Blue 700 Goat anti-mouse IgG (Bio-rad). The proteins were visualized by fluorescence using ChemiDoc MP Imaging Systems (Bio-Rad).

### Immunofluorescence microscopy

Cells were seeded on 24-well plates containing glass coverslips coated with poly-lysine solution (100.000 cells per well). Cells were fixed with 4% PFA in PBS for 15 min, washed twice in PBS, and then permeabilized for 10 min with 0.1% Triton X-100 in PBS. Primary antibodies incubation was carried out for 1h in PBS containing 3% BSA and 0.1% Triton X-100 at 1:100 dilution. Coverslips were washed three times with PBS before secondary anti-mouse antibodies incubation (1:1000 dilution). The antibodies used for immunofluorescence are shown in Supplementary Table 5. Anti-phalloidin was used as a cytoplasmic marker at 1:200, and DAPI (4’6-diamidino-2-phenylindole) (Molecular Probes) was used as a nuclear marker. Coverslips were mounted in Mowiol 4-88 (Sigma-Aldrich). Images were acquired with a confocal laser microscope Leica TCS SP8 STED 3X.

### RNA isolation and sequencing

WT A549, A549-ORF7a and A549-ORF7b cells were seeded (3×10E5) in 6-well plates and lysed using RLT buffer for RNA isolation (RNeasy mini kit, Qiagen). Each sample was performed in triplicate. RNA was isolated following manufacturer protocol, quantified by Nanodrop 1000 (Thermo Scientific) and quality controlled by Bioanalyzer (Agilent). All samples sent for sequencing had a RIN (RNA integrity number) over 9.90. cDNA libraries and sequencing were performed by Novogene Europe, using 400 ng of RNA per sample for library preparation. Samples were sequenced in an Illumina platform using a PE150 strategy.

### Gene sets and expression analysis

Sequencing raw data was quality controlled (error rate, GC content distribution) and filtered, removing bad quality and N-containing sequences and adaptors. Clean data were mapped (HISAT2) to reference genome GRCh38.p13, and gene expression was quantified using FPKM (Fragments Per Kilobase of transcript sequence per Millions of base pairs sequenced). Differential expression analysis was performed using DESeq2 R package (Anders and Huber, 2010).

Raw counts were transformed with the vst function in the DESeq2 package (Love et al., 2014) of the R software version 3.6.3 (R Core Team, 2020), and subsequent PCA was performed with the prcomp function. The 500 genes with the highest variance among samples were considered. Finally, the PCA graph was made with Graphad Prism software. All sequencing data sets are available in the NCBI BioProject database under accession number PRJNA841835.

### GO and Pathways Enrichment Analysis

The annotation function of GO analysis is comprised of three categories: BP, CC, and MF. Kyoto Encyclopedia of Genes and Genomes (KEGG) is a database resource for understanding high-level functions and utilities of the genes or proteins (Kanehisa and Goto, 2000; Kanehisa et al., 2012). GO analysis and KEGG pathway enrichment analysis of candidate DEGs were performed using the R package and using NovoSmart Software. An adjusted p-value less than 0.05 was considered as the cut-off criterion for both GO analysis and pathway enrichment analysis.

### Pathway Enrichment Analysis, Network and PPI Module Reconstruction

To perform the pathway enrichment analysis and the gene network reconstruction, we used the online Metascape tool (http://metascape.org)(Zhou et al., 2019) with the default parameters set. Enrichment analyses were carried out by selecting the genomics sources: KEGG Pathway, GO Biological Processes, Reactome Gene Sets, Canonical Pathways, and CORUM. Terms with p < 0.01, minimum count 3, and enrichment factor >1.5 were collected and grouped into clusters based on their membership similarities. p values were calculated based on accumulative hypergeometric distribution, and q values were calculated using the Benjamini-Hochberg procedure to account for multiple testing. To further capture the relationship among terms, a subset of enriched terms was selected and rendered as a network plot, where terms with similarity >0.3 are connected by edges. Based on PPI enrichment analysis, we ran a module network reconstruction based on the selected genomics databases. The resulting network was constructed containing the subset of proteins that form physical interactions with at least one other list member. Subsequently, employing Molecular Complex Detection (MCODE) algorithm, we first identified connected network components, then a pathway and process enrichment analysis was applied to each MCODE component independently and the three best-scoring (by *p*-value) terms were retained as the functional description of the resulting modules.

### Real time qPCR analysis

RNA samples (500 ng) were reverse transcribed using qScript™ cDNA synthesis kit (Quanta Biosciences Inc.), following manufacturer’s instructions. Primer sequences are available in Supplementary Table 6. The final 15 µL PCR reaction included 2 μL of 1:5 diluted cDNA as template, 3 µL of 5x PyroTaq EvaGreen qPCR Mix Plus with ROX (Cultek Molecular Bioline, Madrid, Spain), and transcript-specific forward and reverse primers at a 10 μM final concentration. Real time PCR was carried out in a QuantStudio 12K Flex system (Applied Biosystems) under the following conditions: 15 min at 95 °C followed by 35 cycles of 30 s at 94 °C, 30 s at 57 °C and 45 s at 72 °C. Melting curve analyses were performed at the end, in order to ensure the specificity of each PCR product. Relative expression results were calculated using GenEx6 Pro software (MultiD-Göteborg, Sweden), based on the Cq values obtained. Statistical differences in expression among groups were assessed using Student’s t-test, setting statistical significance at *P* <0.05.

### Flow cytometric analysis

Flow cytometry was used to analyze the expression levels of ICAM-1. Cells were harvested, washed in PBS and blocked with goat serum for 20 minutes at 4°C. For staining, cells were resuspended in PBS 0.1% BSA 0.01% NaN3 containing the monoclonal antibody PE mouse anti-human CD54 (BD Pharmingen) or its isotype control at a concentration of 1/500. Both antibodies were previously titrated to determine their concentration. Live/Dead Fixable Aqua dye (Invitrogen) was used at 1/1000 to assess cell viability. Cells were incubated for 30 min at 4°C protected from the light, washed with PBS 0.1% BSA 0.01% NaN3 and resuspended in PBS. For these experiments, a CytoFLEX flow cytometer (Beckman Coulter) was used and data was analyzed using FlowJo v10 (BD Biosciences).

### Cell Adhesion Assays

Cell to ECM adhesion was tested using the CytoSelect™ 48-well Cell Adhesion Assay (CBA-070, Cell Biolabs) according to the manufacturer’s protocol. Briefly, cells were seeded at 0.12 × 10^6^ onto the 48-well plate and were left to incubate for 90 min at 37 °C, with 5% CO_2_. The unbound cells were washed off and the adherent cells were stained for 10 min at RT. Any residual stain was removed with 5 washes using dH_2_O and the plate was left to air dry. Using an orbital shaker, the plate was incubated for a final 10 min at RT with extraction solution. The optical density (OD) was read at 595 nm. Bovine serum albumin (BSA) was used as a negative control.

The cell-cell adhesion assay was performed as follow. Briefly, 5 × 10^4^ A549 cells were seeded into a 96-well plate for 24 hours. After 24 h incubation, A549 cells were labeled with calcein-AM. They were co-cultured with the monolayer of A549 cells for 2 h at 37 °C, 5% CO_2_. After the indicated period, non-adherent cells were removed, and then calcein-AM measured the fluorescence using a fluorescein filter set (absorbance maximum of 480/20 nm and an emission maximum of 530/25 nm) to calculate the number of adherent cells.

### ZO-1 staining

For the optical characterization of tight junctions, cells were grown on Transwell membranes (ref. 3401, Costar, 0.4 μm pore size). Seeding density was 70□000 cells. ZO-1 staining was performed after 8 days as described below. Cells were washed three times with PBS and fixed with -20°C methanol for 5 minutes at 4°C. Afterward, the samples were permeabilized with 0.1% Triton X-100 for 5 minutes at RT. Subsequently, a blocking step with PBS containing 1% BSA was performed. The primary anti-ZO-1 antibody (ref. 61-7300, Invitrogen) was diluted at 1:100 in PBS containing 1% BSA and incubated for 3 h at RT. The secondary antibody (anti-rabbit IgG-FITC, ref. 9887, Sigma) was diluted 1:200 in PBS and incubated for 1 h at RT. Cell nuclei were counterstained with DAPI (0,5 μg/mL) for 2 min at RT. Transwell membranes were then sliced into strips and mounted onto glass slides with the cell side up, and treated with anti-fade reagent overnight at 4°C with coverslips on top. Samples were analyzed by a confocal laser scanning microscopy.

## Acknowledgments

The authors wish to acknowledge Bioinformatics & Biostatistics Service at CIB and Dr Aurora Gómez-Durán. This research work was funded by: Junta de Andalucía and the European Commission – NextGenerationEU (Regulation EU 2020/2094), through CSIC’s Global Health Platform (PTI Salud Global).

## Author contributions

Conceptualization: MM and JJG; Methodology: TGG, RFR, NR, SZL, BDLA, JMSC and AJM; Investigation: TGG, RFR, NR, SZL, BDLA, JMSC and AJM; Writing Original Draft: TGG; Writing-Review & Editing: MM and JJG; Supervision: MM and JJG; Project administration: MM and JJG; Funding acquisition: MM and JJG.

## Competing interests

Authors declare that they have no competing interests.

## Notes

### Competing Interest Statement

The authors have declared no competing interest.

