## Supplementary Figures for "Antiviral immune responses, cellular metabolism and adhesion are differentially modulated by SARS-CoV-2 ORF7a or ORF7b"

### Figure S1

**a**

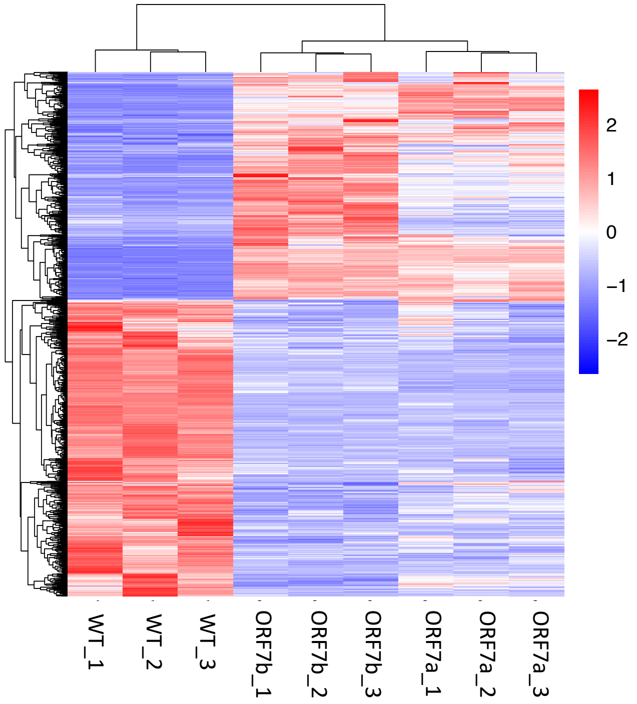


**b**

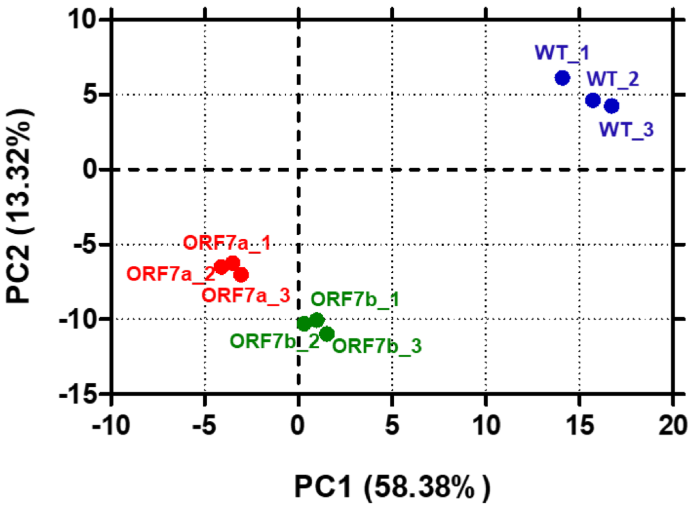


**Figure S1. RNA-Seq analysis of A549 cells expressing SARS-CoV-2 ORF7a or SARS-CoV-2 ORF7b. a,** Heatmap of RNA-Seq data indicating the DEGs patterns between A549 WT (n=3), ORF7a (n = 3) and ORF7b (n = 3). **b,** PCA of A549-WT and transduced A549 cells with ORF7a or ORF7b.

### Figure S2

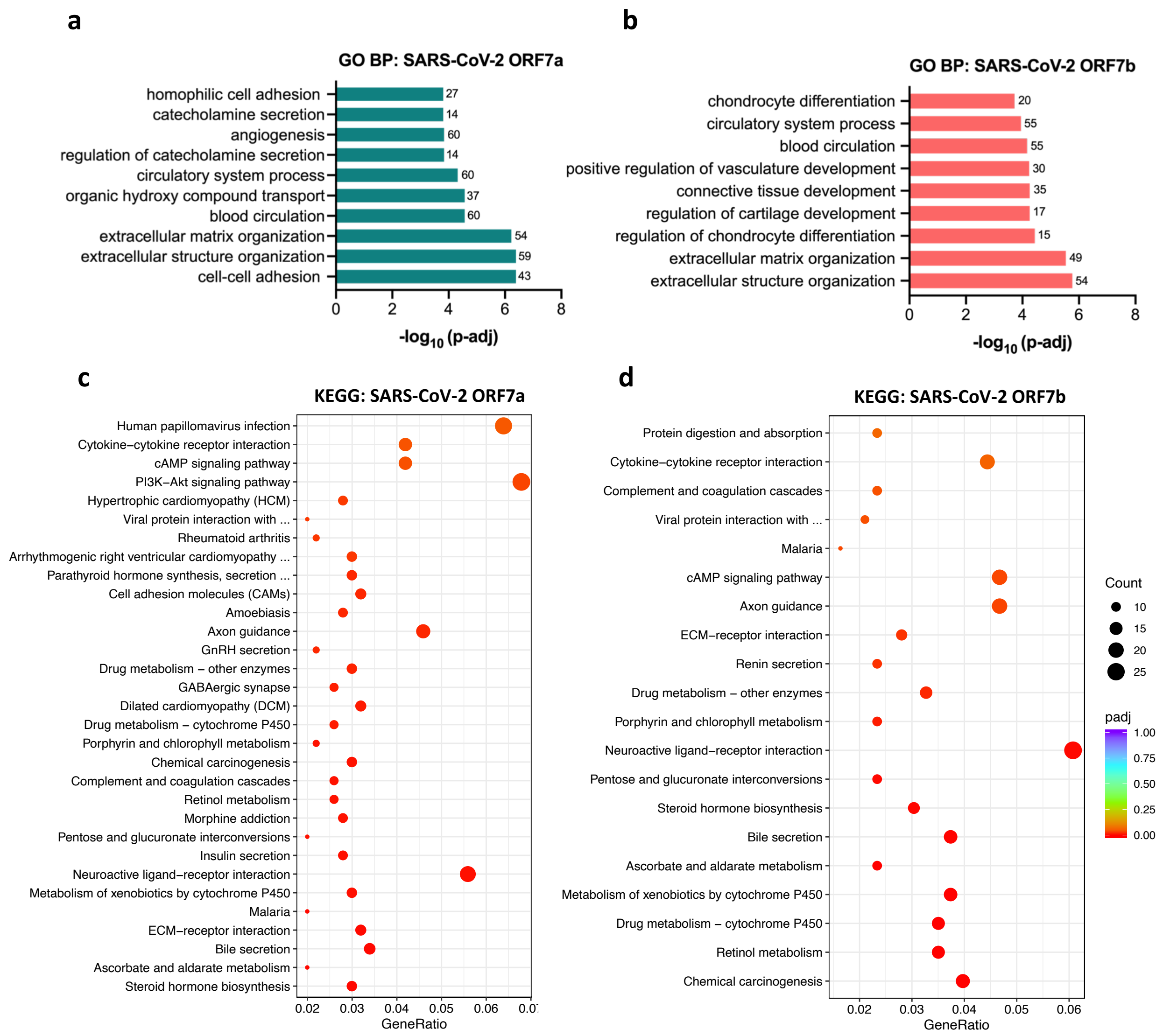


**Figure S2. Enrichment analysis of A549 cells expressing SARS-CoV-2 ORF7a or SARS-CoV-2 ORF7b. a-b,** RNA-Seq data reveals GO enrichment with the top ten classification of biological process in A549 cells expressing ORF7a (**a**) and A549 cells expressing ORF7b compared with A549-WT cells (**b**). **c-d,** Dotplot visualization of enriched KEGG term in ORF7a (**c**) and ORF7b cells (**d**). The color of the dots represent the adjusted p value for each enriched term, and size represents the number of hit genes found in the datasets.

### Figure S3

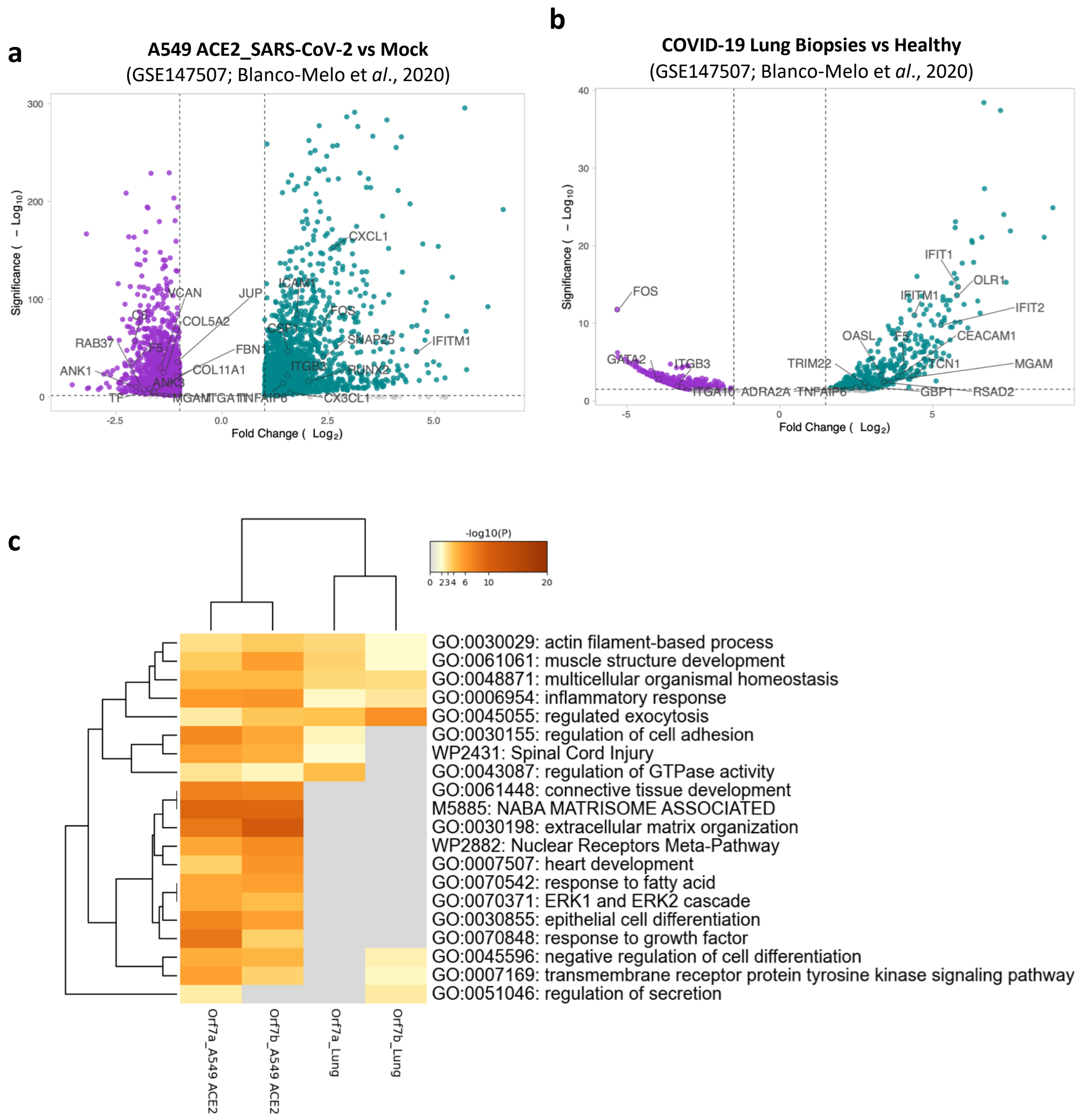


**Figure S3. a-b**, Volcano plot of DEGs in A549 cells transduced with an ACE2 vector and infected with SARS-CoV-2 (a) or post-mortem lung biopsies of COVID-19 patients compared with healthy lung (b). Data are from Blanco-Melo et *al.,* 2020. ORF7a- and ORF7b-associated genes are highlighted. **c**, Heatmap of top 20 enriched ontology clusters across the SARS-CoV-2 RNAseq from Blanco-Melo et *al.,* 2020 and this study, colored by *p*-values (metascape.org).
