## Supplementary Tables S5-S6 for "Antiviral immune responses, cellular metabolism and adhesion are differentially modulated by SARS-CoV-2 ORF7a or ORF7b"

### Supplementary Table 5. Antibody list

| Target protein | Supplier | Catalog number |
|---|---|---|
| Mouse Anti-StrepTag | Sigma-Aldrich | SAB2702215 |
| Mouse Anti-GAPDH | GenScript | A00084 |
| GM130 | abcam | ab52649 |
| Tom20 | abcam | ab186735 |
| Rab4 | abcam | ab109009 |
| Rab7 | abcam | ab126712 |
| PE mouse anti-human CD54 | BD Pharmingen | 560971 |
| PE Mouse IgG1, K Isotype control | BD Pharmingen | 550617 |
| Mouse Anti-$\gamma$-catenin (A-6) | Santa Cruz Biotechnology | sc-514115 |
| Rabbit Anti-ZO-1 | Invitrogen | 61-7300 |
| StarBright Blue 700 Anti-mouse | Biorad | 12004159 |
| Anti-Rabbit IgG-FITC | Sigma | 9887 |

### Supplementary Table 6. Primer list

| Target gen | Forward primer (5'-3') | Reverse primer (5'-3') | Reference |
|---|---|---|---|
| UGT1A9 | GAGGAACATTTATTATGCCACCG | TGCCCAAAGCATCAGCAATT | (1) |
| CYP1A1 | CCTTCGTCCCCTTCACCAT | ATGGTTGATCTGCCACTGGTTT | This work |
| PTGS2 | GAATCATTCACCAGGCAAATTG | CATAAAGCGTTTGCGGTACTCA | This work |
| OASL | TCCAGAACTCATCCTGTCAATCA | GGCCTGGGATAACTCATTGTAAA | This work |
| IFIT1 | TTCAATGAGCCAAAGTCAAGGT | CAGGCAGCCTCTGGAAGCTGGA | This work |
| IFIT2 | GTCCAGAGCTTGACTGTGAGGAA | GAGCCTTCTCAAAGCACACCTT | This work |
| IL8 | GAAGAAACCACCGGAAGGAA | CAAAACTGCACCTTCACACA | This work |

| | | | |
|---|---|---|---|
| IL11 | GAAACAGCAGGCTACAAAACCACT | ACCTCTCTCCTTTGACCTGGAGAC | This work |
| CXCL1 | GGAGCCGTTGGTCAGAAATA | CCTACTGGGCCTCAAATGAA | This work |
| ICAM1 | CGGCTGACGTGTGCAGTAATAC | TGGCTTCGTCAGAATCACGTT | This work |
| ITGB2 | CATTCTCCTGCTGGTCAT | CTTCACCAAGTGCTCCTAAC | This work |
| GAPDH | TGGGTGTGAACCATGAGAAG | TGGCAGTGATGGCATGGAC | This work |

1. Ramírez, J., Mirkov, S., Zhang, W., Chen, P., Das, S., Liu, W., Ratain, M. J., and Innocenti, F. (2008) Hepatocyte nuclear factor-1 alpha is associated with UGT1A1, UGT1A9 and UGT2B7 mRNA expression in human liver. *Pharmacogenomics J.* 8, 152–161
